# eIF4A controls translation of estrogen receptor alpha and is a therapeutic target in advanced breast cancer

**DOI:** 10.1101/2024.05.08.593195

**Authors:** Jacob A. Boyer, Malvika Sharma, Madeline A. Dorso, Nicholas Mai, Corina Amor, Jason M. Reiter, Ram Kannan, Sunyana Gadal, Jianing Xu, Matthew Miele, Zhuoning Li, Xiaoping Chen, Qing Chang, Fresia Pareja, Stephan Worland, Douglas Warner, Sam Sperry, Gary G. Chiang, Peggy A. Thompson, Guangli Yang, Ouathek Ouerfelli, Elisa de Stanchina, Hans-Guido Wendel, Ezra Y. Rosen, Sarat Chandarlapaty, Neal Rosen

## Abstract

The majority of human breast cancers are dependent on hormone-stimulated estrogen receptor alpha (ER) and are sensitive to its inhibition. Treatment resistance arises in most advanced cancers due to genetic alterations that promote ligand independent activation of ER itself or ER target genes. Whereas re-targeting of the ER ligand binding domain (LBD) with newer ER antagonists can work in some cases, these drugs are largely ineffective in many genetic backgrounds including ER fusions that lose the LBD or in cancers that hyperactivate ER targets. By identifying the mechanism of ER translation, we herein present an alternative strategy to target ER and difficult to treat ER variants. We find that ER translation is cap-independent and mTOR inhibitor insensitive, but dependent on 5’ UTR elements and sensitive to pharmacologic inhibition of the translation initiation factor eIF4A, an mRNA helicase. EIF4A inhibition rapidly reduces expression of ER and short-lived targets of ER such as cyclin D1 and other components of the cyclin D-CDK complex in breast cancer cells. These effects translate into suppression of growth of a variety of ligand-independent breast cancer models including those driven by ER fusion proteins that lack the ligand binding site. The efficacy of eIF4A inhibition is enhanced when it is combined with fulvestrant—an ER degrader. Concomitant inhibition of ER synthesis and induction of its degradation causes synergistic and durable inhibition of ER expression and tumor growth. The clinical importance of these findings is confirmed by results of an early clinical trial (NCT04092673) of the selective eIF4A inhibitor zotatifin in patients with estrogen receptor positive metastatic breast cancer. Multiple clinical responses have been observed on combination therapy including durable regressions. These data suggest that eIF4A inhibition could be a useful new strategy for treating advanced ER+ breast cancer.

## Introduction

Estrogen receptor alpha (ER) is a ligand activated transcription factor and member of the extended family of nuclear receptors^1,2^. Upon estrogen binding, ER translocates to the nucleus, dimerizes, and induces the transcription of an ensemble of genes involved in proliferation, lineage specification, and other context specific functions^3^. ER is required for development of the mammary ductal epithelium and 70-75% of breast tumors retain dependence on ER for growth/division ^4–6^. Hormonal therapies targeting ER are highly active in these ER+ metastatic breast cancers and have been remarkably successful in improving outcomes. Unfortunately, resistance to hormonal therapy is nearly universal, and over 90% of patients develop resistance to various drugs targeting ER^7^.

Dysregulation of the PI3K/AKT/mTOR pathway is common in ER-dependent breast cancer, and activating mutations of *PIK3CA*, the catalytic subunit of class 1 PI3 Kinase, occur in 40% of these tumors^8^. The PI3K pathway enhances the proliferation, motility and invasiveness of these tumor cells and activation of the mTOR complex 1 (mTORC1) maintains high levels of eIF4E-dependent protein translation^9,10^ ^11–14^. Clinically, combination PI3K/ER inhibition is an effective strategy but efficacy is partly diminished by enhanced ER signaling and ER expression following PI3K inhibition^15,16^. Inhibition of mTORC1 decreases total protein translation by as much as 70%^17^. However, for a select number of short lived-proteins, expression can be maintained using non-canonical mechanisms of protein translation^18–23^. Prior studies have estimated that ER is a short-lived protein with a half-life of approximately 3-6 hours ^24–26^. We therefore asked how despite being a short-lived protein, ER expression is maintained when PI3K/mTOR is inhibited.

Here we show that, during mTOR inhibition, continued expression of ER is maintained by eIF4E-independent translation. This is dependent upon the RNA helicase, eIF4A, a component of the eIF4F initiation complex and elements of the 5’ untranslated region (5’ UTR) of the ER-encoding mRNA, *ESR1.* Pharmacological inhibition of eIF4A reduces the expression of ER as well as a number of other short half-life proteins controlling cell cycle entry including cyclin D1, cyclin D3 and CDK4. EIF4A inhibitors effectively reduce ER expression in hormone receptor dependent tumor models driven by wild type ER, hormone-insensitive ER mutants or ER fusion proteins. Moreover, inhibition of ER translation via eIF4A blockade combined with induction of ER degradation by the ER degrader fulvestrant causes synergistic inhibition of ER expression and inhibition of breast tumor growth in xenograft models. These data suggest that combined inhibition of eIF4A with fulvestrant could be a novel strategy for the treatment of breast cancers with acquired resistance to hormone receptor inhibition. We therefore initiated a phase I/II clinical trial of the combination in such patients. The trial (NCT04092673) employs the eIF4A inhibitor zotatafin, in combination with fulvestrant. Data from this trial shows that this combination is well tolerated^27^ and multiple tumor regressions have been observed in heavily pre-treated endocrine therapy resistant patients.

## Results

### Estrogen receptor alpha expression is eIF4A dependent

Canonical eukaryotic translation is initiated by eukaryotic initiation factor 4E (eIF4E), which upon binding the m7G mRNA cap, nucleates an initiation complex comprised of eIF4E, eIF4G, and eIF4A, all together known as eIF4F^28,29^. Multiple oncogenic pathways, most notably mTORC1, control translation initiation by phosphorylating and sequestering 4EBP1 away from eIF4E^17,30^. Once assembled, the immature ribosome begins “scanning” within the 5’UTR, and this process is aided by the RNA helicase and DEAD box containing protein eIF4A, which facilitates unwinding of complex structures in the 5’ UTR^31–33^ (Figure 1A). By activating 4EBP1, mTOR inhibitors reduce global translation, but some mRNA transcripts are insensitive to mTOR inactivation^17,34–36^ Given the previous work showing that ER activity and expression is enhanced when PI3K/mTOR is inhibited^15,16^, we hypothesized that ER might be one such eIF4E-independently translated protein. To test this, we treated ER+ breast cancer cell line MCF7 with the potent mTORC1/2 inhibitor, RapaLink-1 ^37^ (Figure 1B). mTOR effectors, AKT, S6 and 4EBP1 were all dephosphorylated by four hours post treatment, and this inhibition was accompanied by a reduction in global protein synthesis by up to 75% (Figure 1B, Figure S1A).

**Figure 1:**
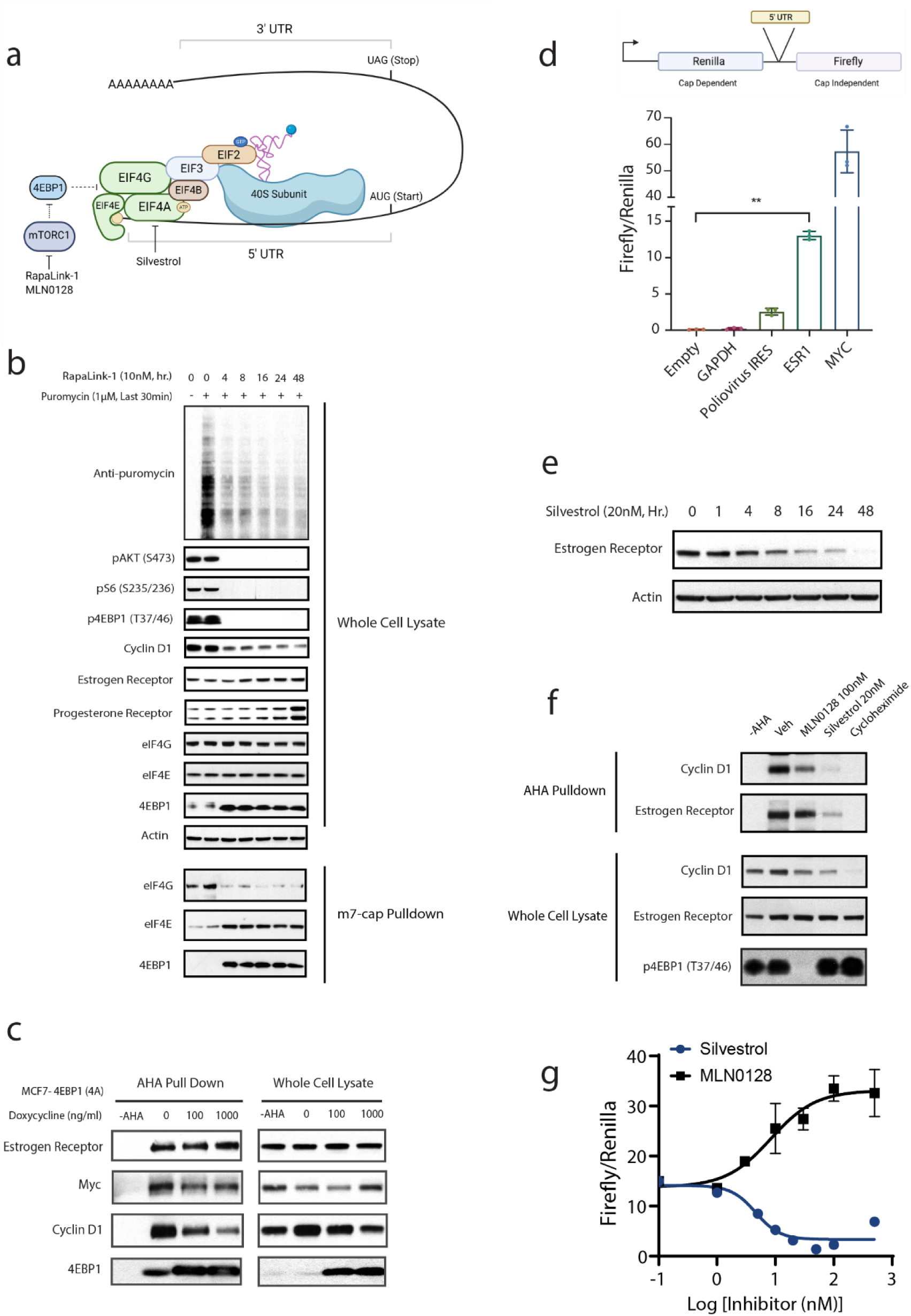
Estrogen Receptor Alpha Expression is EIF4A Dependent (A) Schematic of translation initiation. The EIF4F complex is depicted in green and consists of EIF4E, EIF4G and EIF4A. Inhibitors of EIF4A (Silvestrol) and mTORC1 (RapaLink-1 and MLN0128) are shown. (B) MCF7 cells were treated with 10nM RapaLink-1 for the indicated times. To measure global translation, 1 µM puromycin was pulsed for the last 30min, and incorporation was assessed using an anti-puromycin antibody. For m7G cap pulldowns, 200µg of cell lysates were first incubated for 2hr with m7-Guanosine conjugated agarose beads (Jena Biosciences), washed, eluted and used for immunoblotting. (C) MCF7 expressing doxycycline inducible 4EBP1 (T37A,T46A,S65A,T70A) were plated in 1μg/ml doxycycline for 24hr followed by methionine starvation for 30min. Cells were then pulsed with 100µM L-Azidohomoalanine (AHA) for 2hr. 200µg of lysate were subjected to a click chemistry reaction using biotin conjugated alkyne. AHA labeled proteins were then isolated via streptavidin-agarose assisted precipitation. (D) 1.5 million MCF7 cells were plated in 6cm dishes, and transfected with 2µg of the indicated dual luciferase constructs for 24hr. Firefly and Renilla luciferase activity was measured via luminescence and “cap-independent activity” of the various 5’ UTR elements were quantified as the ratio of Firefly to Renilla. P-values were determined by ordinary one-way ANOVA. P< .01 depicted as ** (E) MCF7 cells were treated with 20nM Silvestrol for the indicated times. (F) MCF7 cells were starved of methionine for 30min in the presence of mTOR inhibitor MLN0128 (100nM), silvestrol (20nM) or cycloheximide (50μg/ml). Cells were then labeled with 100μM L-Azidohomoalanine (AHA) for 2hr. 200 μg of lysate was used for click chemistry with biotin-alkyne or set aside for immunoblotting (Whole Cell Lysate). AHA labeled proteins were then isolated via streptavidin-agarose assisted precipitation. (G) 1.5 million MCF7 cells were plated in 6cm dishes, and transfected with 2µg of the indicated dual luciferase constructs for 24hr. At the same time, cells were treated with increasing doses of INK0128 or silvestrol. Firefly and Renilla luciferase activity was measured via luminescence and “cap-independent activity” of the *ESR1* 5’ UTR construct was quantified as the ratio of Firefly to Renilla. All experiments in this figure were repeated three times, independently.

Dephosphorylation of 4EBP1 coincided with enhanced binding of 4EBP1, and decreased binding of eIF4G to the eIF4E-m7G cap complex (Fig 1B). Dephosphorylation of 4EBP1 was also associated with a reduction in the expression of cyclin D1, translation of which is known to be mTOR/eIF4E dependent ^13,38–40^(Figure 1B). In contrast to cyclin D1, estrogen receptor levels were unchanged during this suppression of both cap-dependent and global translation (Figure 1B). As previously observed, inhibition of AKT was associated with an increase in the ER target, progesterone receptor by 24 hours^15,16^ (Figure 1B). In contrast to mTOR inhibition, blocking all mechanisms of protein translation using cycloheximide resulted in a time dependent decrease of ER expression, with a half-life between 4 and 8 hours (Figure S1B). MTOR controls translation through multiple substrates, including: LARP1, S6K and 4EBP1 ^17,41,42^ but only 4EBP1 engages cap-binding protein, eIF4E, directly. To confirm that ER expression was insensitive to dephosphorylated 4EBP, and is thus translated in an eIF4E independent fashion, we expressed a doxycycline inducible 4EBP1 mutant termed 4EBP1 “4A”, in which the four major sites of mTOR phosphorylation have all been mutated to alanine (T37A/T46A/S65A/T70A)^17,43^. Expression of this mutant ablated cap-dependent translation in a dose dependent manner and induced 4EBP1-eIF4E binding at the expense of eIF4G (Figure S1C). To assess protein translation directly, we used the methionine analog, L-Azidohomoalanine (AHA), to label and isolate *de novo* synthesized proteins^44^(Figure 1C). Translation of cyclin D1 was inhibited as a function of doxycycline dose (4EBP1-4A expression) (Figure 1C), but ER translation was unaffected (Figure 1C). Myc, the translation of which is known to be eIF4E-independent, was similarly unaffected by 4EBP1-4A expression ^19,45,46^(Figure 1C).

Organisms from viruses to mammals have evolved a variety of mechanisms for the eIF4E independent (cap independent) translation of select mRNAs^19,22,23,47–49^ Often the elements facilitating cap-independent translation are contained in the mRNA 5’ untranslated region (5’ UTR) and are termed IRES elements (Internal ribosome entry sites). To test whether the 5’ UTR of ER (*ESR1*) drives cap independent translation, we used a bicistronic luciferase vector^50^ containing a cap-dependently translated renilla luciferase and a firefly luciferase whose cap-independent expression is under control of a chosen insert (Figure 1D). We observed that the 5’ UTR of *ESR1* was capable of driving cap-independent translation approximately 120-fold higher than the empty insert and 5-fold higher than the poliovirus IRES control, but lower than that of a powerful IRES from the *Myc* 5’ UTR (Figure 1D). We additionally confirmed that the cap-independent activity of the *ESR1* 5’ UTR was not an artifact of cryptic promoter activity or read through by using luciferase assay constructs containing the *ESR1* 5’ UTR with a hairpin and lacking a promoter (Figure S1D). These results reveal that while ER is a short-lived protein, its translation can be sustained in an eIF4E independent manner during mTOR inhibition, and this eIF4E independent activity is mediated through elements in the 5’UTR.

IRES elements in the mRNA 5’UTR often have complex secondary structures that bind to a subset of eukaryotic initiation factors that recruit the ribosome. Such structures require remodeling or unwinding by RNA helicases during initiation^47,51^. We hypothesized that eIF4A, the major RNA helicase employed in translation initiation, might control ER protein synthesis. To test this we treated MCF7 with 20nM of the selective^52^ eIF4A inhibitor silvestrol for 48 hours and observed that levels of ER protein declined monotonically throughout the time course, with detectable reductions after 4 hours of drug exposure (Figure 1E). A similar decline in ER protein cooccurred in other models, including ER+ breast cancer cell lines T47D, ZR-75-1 and BT474, as well as the ovarian cancer cell line SKOV3, with ER expression decreasing between 4 and 8 hours following silvestrol treatment (Figure S1E). Silvestrol and its synthetic rocaglate analog CR-31-B (+/-)^53^, both inhibited ER expression at concentrations between 20-30nM, and cyclin D1 expression was reduced at similar doses (Figure S1F). Two other mechanistically distinct inhibitors of eIF4A, pateamine A and hippuristanol^54–56^ also inhibited both ER and cyclin D1 expression by 24 hours (Figure S1G). To assess whether eIF4A controls ER translation directly, we again used AHA to measure *de novo* protein synthesis^44^. We used the fast-acting ATP competitive mTOR inhibitor MLN0128^12^ to block EIF4E dependent translation, silvestrol to inhibit eIF4A, and cycloheximide as a control to block global translation (Figure 1F). The translation of cyclin D1 but not of ER was suppressed by mTOR inhibition, whereas the translation of both cyclin D1 and ER were reduced in the presence of either silvestrol or cycloheximide (Figure 1F). *ESR1* mRNA levels were largely stable during the first 24 hours post silvestrol treatment, decreasing at most by 25% during the first 16 hours. Protein expression however was reduced to 50% of its initial level at 16 hours, implying a non-transcriptional mechanism of ER regulation by eIF4A (Figure S1H.) Finally, we tested whether eIF4A inhibition attenuated the cap-independent translation conferred by the *ESR1* 5’ UTR. As a function of dose, silvestrol effectively blocked the firefly (cap-independent) translation mediated by the 5’ UTR of *ESR1* (Figure 1G). In contrast, the mTOR inhibitor MLN0128 actually increased the cap-independent translation conferred by the *ESR1* 5’ UTR (Figure 1G). Taken together, these results establish that ER is translated in an eIF4A dependent but eIF4E independent manner, and that cyclin D1 translation depends on both eIF4E and eIF4A.

### eIF4A regulates ER activity and cell growth

We analyzed how eIF4A inhibition affected ER function and cell growth. We treated MCF7 for 24 hours with 20nM Silvestrol and observed marked downregulation of the mRNAs of five positively regulated ER target genes (*PGR, GREB1, TFF1, IGFBP4* and *SERPINA1*^57^(Figure 2A). Moreover, silvestrol enhanced the mRNA expression of *TP3INP1*, a gene that is suppressed by ER ^57^(Figure 2A). Consistent with these findings, eIF4A inhibition was also found to block estradiol stimulated gene expression in both MCF7 and T47D models(Figure 2B, Figure S2A). To rule out ER independent effects of silvestrol on ER target gene expression, we performed chromatin immunoprecipitation of estrogen receptor bound to an estrogen response element (ERE) in target gene *TFF1/pS2*. Silvestrol pretreatment reduced ER occupancy of this response element by approximately 2.5-fold and prevented the estradiol induced ER occupancy at this site by a similar magnitude (Figure 2C). Overall, these data suggest that reducing ER expression by blocking eIF4A results in a decreased breast cancer response to estrogen *in vitro*. We asked whether eIF4A inhibition affected the proliferation of ER+ breast cancer models. ER dependent MCF7 (Figure 2D) and T47D (Figure 2E) cells were treated with increasing doses of silvestrol, and in both models, growth was suppressed in a dose dependent manner, with 5nM blocking growth by 50% at 3 days and 20nM completely blocking growth across 7 days. (Figure 2D, 2E). These results demonstrate that as a single agent, eIF4A inhibitors can suppress ER expression and ER target genes, as well as block growth of ER+ breast cancer models.

**Figure 2:**
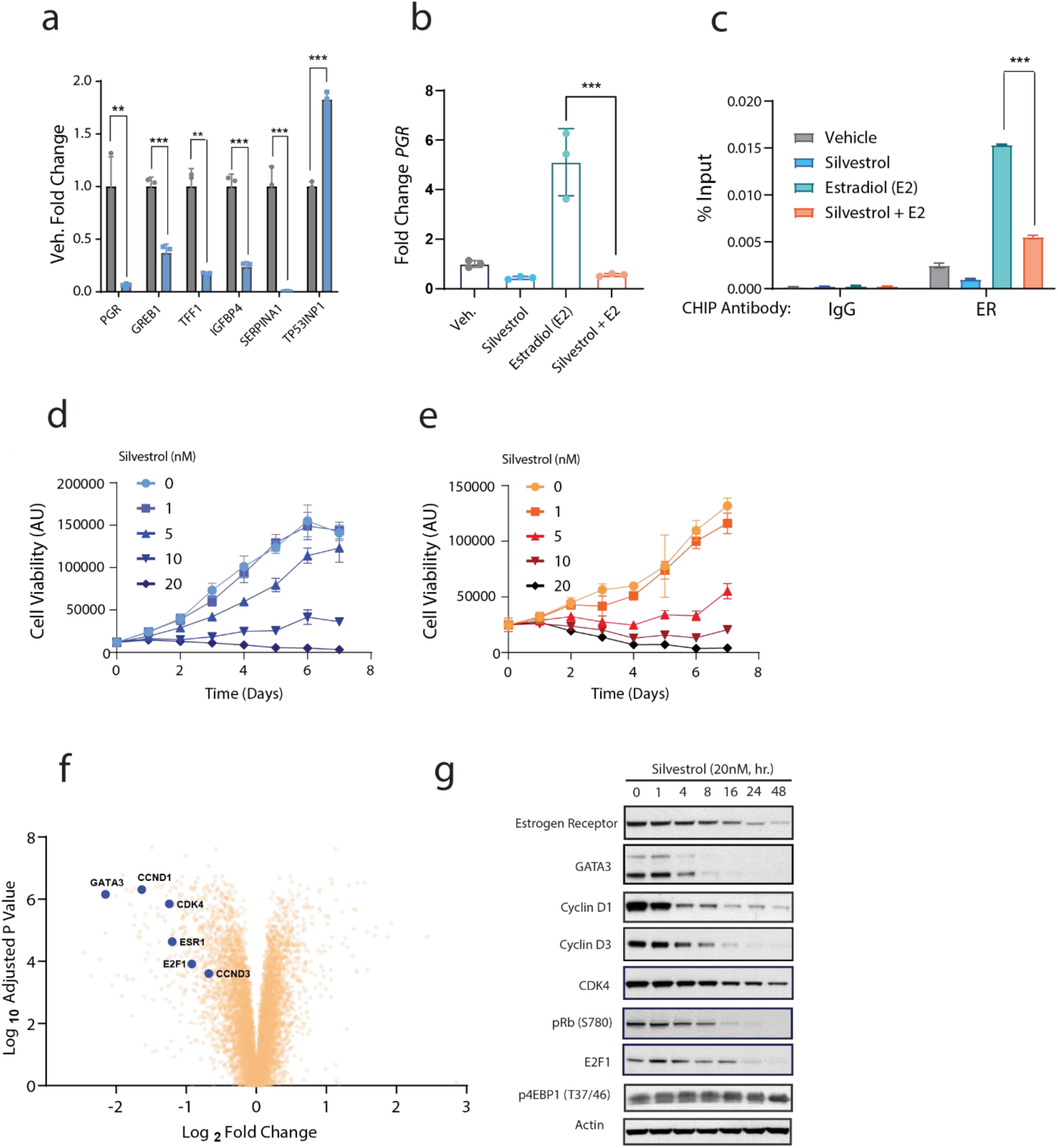
eIF4A regulates ER activity and cell growth (A) MCF7 treated for 24hr with 20nM Silvestrol followed by analysis of canonical ER target gene expression by RT-qPCR. P-values were determined by students t-test for each gene. P< .01 depicted as ** and p< .001 as ***. Data is representative of three independent experiments. (B) MCF7 were plated in DMEM F12 containing charcoal stripped FBS and lacking phenol red, followed by treatment with silvestrol (20nM) for 24hr. Cells were then stimulated with 10nM estradiol for an additional 24hr. *PGR* mRNA expression was analyzed by RT-qPCR. P-values were determined by ordinary one-way ANOVA. P< 0.001 is depicted as ***. Data is representative of three independent experiments. (C) MCF7 were placed in DMEM F12 containing charcoal stripped FBS and lacking phenol red, followed by treatment with 20nM Silvestrol for 24hr. Cells were then stimulated with 10nM estradiol for 1hr. ER binding to the *TFF1* enhancer element was analyzed by Chromatin Immunoprecipitation Assay (ChIP) and quantified by RT-qPCR. Representative data is shown with technical replicates for n=3. P-values were determined by ordinary one-way ANOVA. P< 0.001 is depicted as ***. Data is representative of three independent experiments. (D) MCF7 cells were plated in 96 well plates and treated for up to seven days with increasing doses of silvestrol. Cell viability was measured at each timepoint using ATP-glo luminescence. Data is representative of three independent experiments. (E) T47D cells were plated in 96 well plates and treated for up to seven days with increasing doses of silvestrol Cell viability was measured at each timepoint using ATP-glo luminescence. Data is representative of three independent experiments. (F) MCF7 were treated with Veh. (DMSO) or silvestrol (20nM) in triplicate for 24hr. followed by analysis of protein expression by tandem mass tag LC-MS. Significant hits were those proteins changing by at least log2 fold change>1 and an adj. P-value of <.05 (see methods). (G) MCF7 were treated with 20nM Silvestrol for the indicated times. Data is representative of three independent experiments.

Many transcripts have been described as sensitive to eIF4A inhibition, and eIF4A inhibitors can block the growth of various cancer models^52,58–63^. We therefore sought to determine the global protein changes that occur in ER+ breast cancer models after eIF4A inhibition. MCF7 cells were treated with DMSO or 20nM silvestrol for 24 hours followed by proteomic analysis by tandem mass tag (TMT) LC-MS (Figure 2F). This facilitated the identification of proteins that are synthesized in an eIF4A dependent manner and that have half-lives short enough to detect inhibition of their expression after blocking synthesis within this time frame. 122 proteins were statistically reduced by at least 2-fold after this experiment (Table S1), with several of potential relevance to ER+ breast cancer growth. (Figure 2F). We observed a greater than 2-fold reduction in estrogen receptor alpha (*ESR1*) and a 4-fold reduction in the ER co-factor *GATA3,* two drivers of ER+ breast cancer ^64,65^ (Figure 2F). We also noted statistically significant decreases in multiple cell cycle regulators including CDK4, E2F1, cyclin D3 (*CCND3*), and previously identified eIF4A target cyclin D1 (*CCND1*)^66,67^(Figure 2F). To validate this result, we treated MCF7 cells with 20nM silvestrol for up to 48 hours and analyzed the expression of several of these targets by immunoblotting, finding that ER levels were reduced as previously shown (Figure 2G). GATA3 expression was also inhibited by silvestrol, and expression decreased to a minimum by 8 hours (Figure 2G). Proteins regulating the G1/S checkpoint, including cyclin D1, cyclin D3 and CDK4 all decreased, albeit with varying kinetics (Figure 2G). Cyclin D1 expression was reduced by one hour post treatment and continued to decrease until reaching a minimum at the 8 hour timepoint (Figure 2G). Reductions in G1 cyclins and in CDK4 resulted in attenuated Rb phosphorylation beginning 8 hours post treatment (Figure 2G). Overall, these data suggest that eIF4A inhibition reduces ER expression and suppresses ER dependent transcription. eIF4A inhibition also potently suppresses growth of ER+ breast cancer models, likely through regulation of ER and other factors including G1 cyclins.

### eIF4A inhibition combined with Fulvestrant synergistically reduces ER expression

Given our results showing that eIF4A inhibition suppresses ER and G1 cyclins, as well as cell growth, we wondered whether targeting eIF4A in combination with clinically used anti-endocrine therapies would be an effective anti-tumor strategy. Selective Estrogen Receptor Degraders (SERDs) such as fulvestrant are ER antagonists which not only block receptor signaling but also induce ER degradation^57,68,69^. We reasoned that blocking ER synthesis with an eIF4A inhibitor, while simultaneously enhancing ER turnover with a SERD, would cause synergistic reductions in ER expression. To test this, we treated MCF7 with 30nM fulvestrant, 20nM silvestrol or the combination for up to 24hr. Fulvestrant rapidly reduced ER expression one hour after treatment, reaching its maximal effect at 4 hours (Figure 3A), and suppression was maintained for 16 hours followed by rebound of ER levels at 24 hours (Figure 3A). Silvestrol suppressed ER expression more slowly (Figure 3A), and expression was blocked at 8 hours post-treatment and continued over the next 16 hours (Figure 3A). We quantified the area under the curve for ER expression through time for each drug alone and in combination (Figure S3A). Cumulative ER levels in the presence of fulvestrant and silvestrol were calculated to be 935.7 and 867.2 respectively. In the presence of the combination, cumulative ER levels were calculated to be 215.2, representing a 4 fold better suppression of ER levels using the combination than with either silvestrol or fulvestrant alone (Figure S3A). To quantify the effects of silvestrol and fulvestrant on estradiol induced gene expression, we used T47D cells expressing an ERE driven luciferase cassette^70^ (Figure 3B). The lower doses of either silvestrol (5nM) or fulvestrant (3nM) blocked the ability of estradiol to activate the reporter(Figure 3B). Combining both compounds however not only blocked estradiol stimulated gene expression, but reduced baseline ER dependent gene expression to half of that the control (Figure 3B). Higher doses of either silvestrol (20nM) or fulvestrant (30nM) again suppressed the ability of estradiol to stimulate the reporter, while the combination reduced basal ER driven gene expression to five times lower than the unstimulated control (Figure 3B). Combination treatment with both drugs was twice as effective at suppressing estradiol stimulated PGR expression, and seven times more effective at suppressing *TFF1/pS2* expression than either compound alone (Figure S3B). These results reveal that the combinatorial reduction of ER expression using eIF4A inhibitors and SERDs can lead to major improvements in target pathway inhibition.

**Figure 3:**
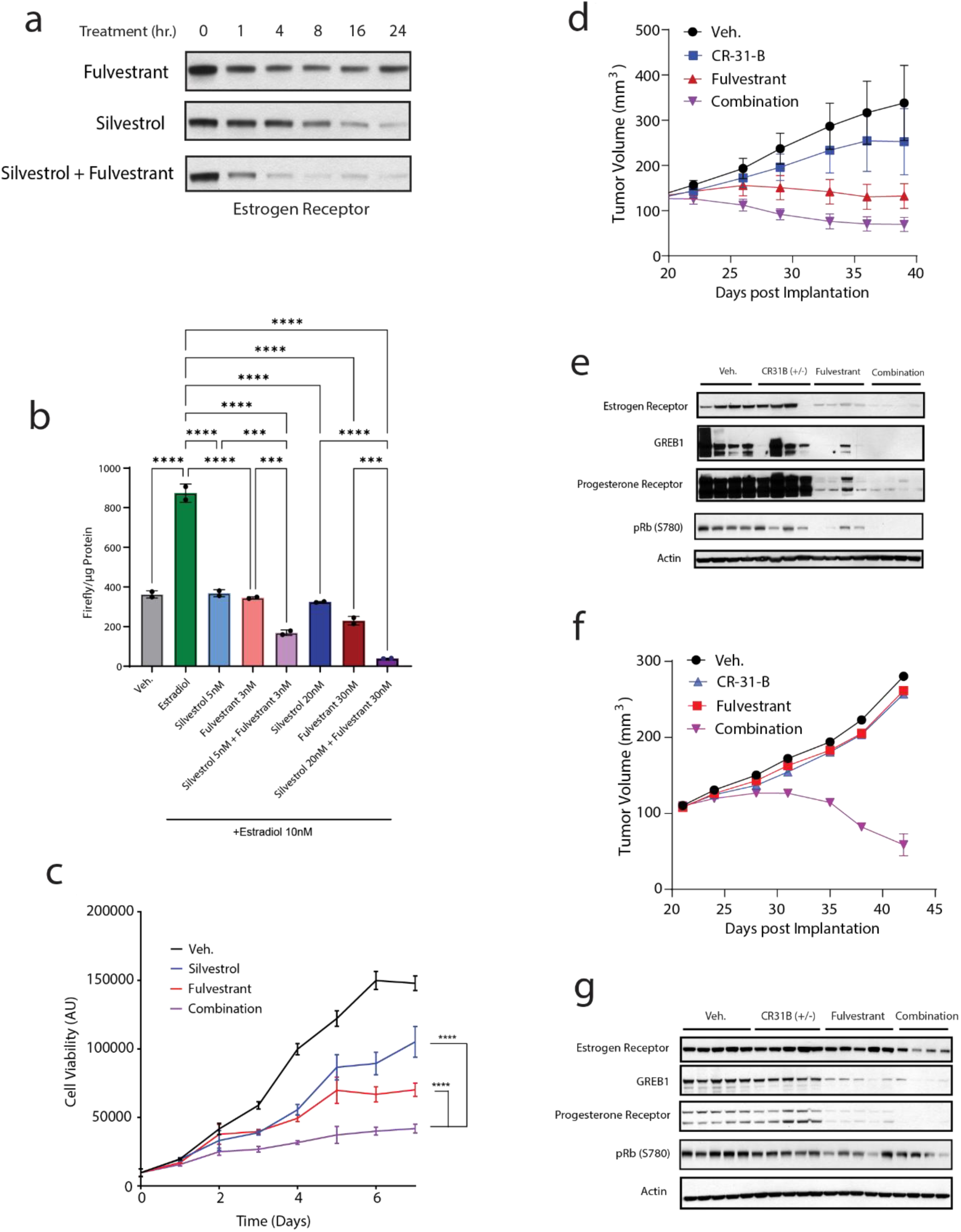
eIF4A inhibition combined with fulvestrant minimizes ER expression and blocks tumor growth (A) MCF7 cells were treated for indicated times with Silvestrol (20nM), Fulvestrant (30nM) or combination. Data is representative of three independent experiments. (B) T47D kBluc were placed in DMEM F12 containing charcoal stripped FBS and lacking phenol red with or without the indicated doses of silvestrol or fulvestrant at the indicated doses for 24hr. Cells were then stimulated with 10nM estradiol for an additional 24hr. Firefly expression was quantified via luminescence and normalized to protein mass obtained via BCA protein quantification. N=2 replicates for each group. P-values were determined by ordinary one-way ANOVA. P< .001 is depicted as ***. P<.0001 is depicted as ****. Data is representative of three independent experiments. (C) MCF7 were treated with either 5nM silvestrol, 3nM fulvestrant or the combination for up to seven days. Cell growth was measured daily via ATP-glo luminescence. P-values were determined by ordinary one-way ANOVA at day seven. P< .0001 depicted as ****. Data is representative of three independent experiments. (D) Nude mice were implanted with estrogen pellets (0.18 mg) for 3d before injection of MCF7 10million cells/mouse. Once tumors reached 100 mm^3^ mice were treated twice weekly with 200mg/kg Fulvestrant administered subcutaneously, 1mg/kg CR-31-B (+/-) administered by i.v. or the combination. Data is representative of two independent experiments. (E) Immunoblots from xenografts in Fig. 3D collected 24 hours following the final dose of the indicated compounds. (F) Nude mice were implanted with estrogen pellets (0.72 mg) for 3 days before injection of MCF7 10million cells/mouse. Once tumors reached 100 mm^3^ mice were treated twice weekly with 200mg/kg Fulvestrant administered subcutaneously, 1mg/kg CR-31-B (+/-) administered by i.v. or the combination. Data is representative of two independent experiments. (G) Immunoblots from xenografts in figure 3F collected 24hr following the final dose of the indicated compounds.

To determine if these differences translated to improved effects on tumor growth, we treated MCF7 cells with silvestrol, fulvestrant, or the combination for up to 5 days. Each compound was observed to block cell growth by approximately 50% at day 3 (Figure 3C). The combination inhibited cell growth at least twice as well as either compound alone(Figure 3C). In addition to ER, eIF4A inhibition reduced the expression of several cell cycle mediators including cyclin D1. We therefore investigated the effects of combined eIF4A inhibitor and fulvestrant treatment on cell cycle progression. Using EdU labeling, we examined the percentage of cells in the S phase following treatment with silvestrol, fulvestrant or the combination for 48 hours. At baseline, approx. 30% of MCF7 cells were in S phase(Figure S3C). Single agent silvestrol significantly lowered this fraction to 7.5%, while fulvestrant lowered the S phase fraction to approx. 20%(Figure S3C). The combination of fulvestrant with silvestrol was highly effective at reducing the fraction of S phase cells; lowering the percentage of S phase cells to 0.5% (Figure S3C).

To test the effects of these inhibitors *in vivo*, we treated MCF7 xenografts with the eIF4A inhibitor, CR-31-B ^63^, fulvestrant or the combination. To determine if estrogen levels affected the anti-tumor efficacy of this combination, we used mice implanted with either low (0.18mg) or high (0.72mg) estrogen pellets (Figure 3D-G). In the low estrogen context, fulvestrant was sufficient to inhibit the growth of the xenografts over a 20 day period, while CR-31-B had little to no effect in this time frame (Figure 3D). The combination of CR-31-B and fulvestrant had a greater antitumor effect than fulvestrant alone—preventing tumor growth and inducing a mild regression (Figure 3D). In the high estrogen setting, neither fulvestrant nor CR-31-B had a substantial effect when given alone. The combination however produced a profound, durable regression lasting 45 days (Figure 3F). In both cases, the combination treatment suppressed the expression ER and ER targets, progesterone receptor and GREB1, better than either compound alone (Figure 3E and 3G). Additionally, in the low estrogen setting, the combination treatment more potently suppressed Rb phosphorylation, suggesting better suppression of cell cycle entry when both drugs are given (Figure 3E). These drugs were well tolerated alone or in combination, and mice did not exhibit weight loss over the course of the study (Figure S3D). Taken together, these data suggest that combinations of eIF4A inhibitors with ER degraders such as fulvestrant, reduce ER expression better than either compound alone, and this combination may be an effective strategy for treating ER+ breast cancers.

### eIF4A inhibition blocks the expression of clinically significant ER variants

Resistance to anti-estrogen therapy often arises due to ER somatic mutations which restore estrogen receptor signaling^71–79^. One such example is the ER-D538G mutant which is the most frequent ESR1 mutation detected^80^. This mutation allows a relaxed conformation of helix-12, mimicking an agonist bound receptor confirmation, thus conferring reduced hormone dependence, hyperactivity, and resistance to anti hormonal therapy^73–75^. We used a previously characterized MCF7 derived cell line expressing a knocked in version of ESR1-D538G, which retains the endogenous *ESR1* untranslated elements^74,75^. CR-31-B treatment reduced the levels of both D538G ER and cyclin D1 in a dose dependent manner, with 30nM producing saturable inhibition of both targets (Figure 4A). Silvestrol treatment produced a similar effect and 20nM suppressed both ER and cyclin D1 expression in both wildtype and D538G ER expressing cells (Figure S4A). The stability of estrogen receptor D538G was similar to that of wildtype ER, and expression was notably reduced 4 and 8 hours after treatment (Figure S4B). As previously shown, cells expressing ER D538G were less sensitive to fulvestrant^74^, (IC50 of 5.4nM compared to 0.41nM for the wildtype (Figure 4B). In contrast, sensitivity of wild type and ER-D538G cells to CR-31-B was similar in the the wildtype (3.9nM) and ER-D538G expressing cells (4.1nM) (Figure 4C).

**Figure 4:**
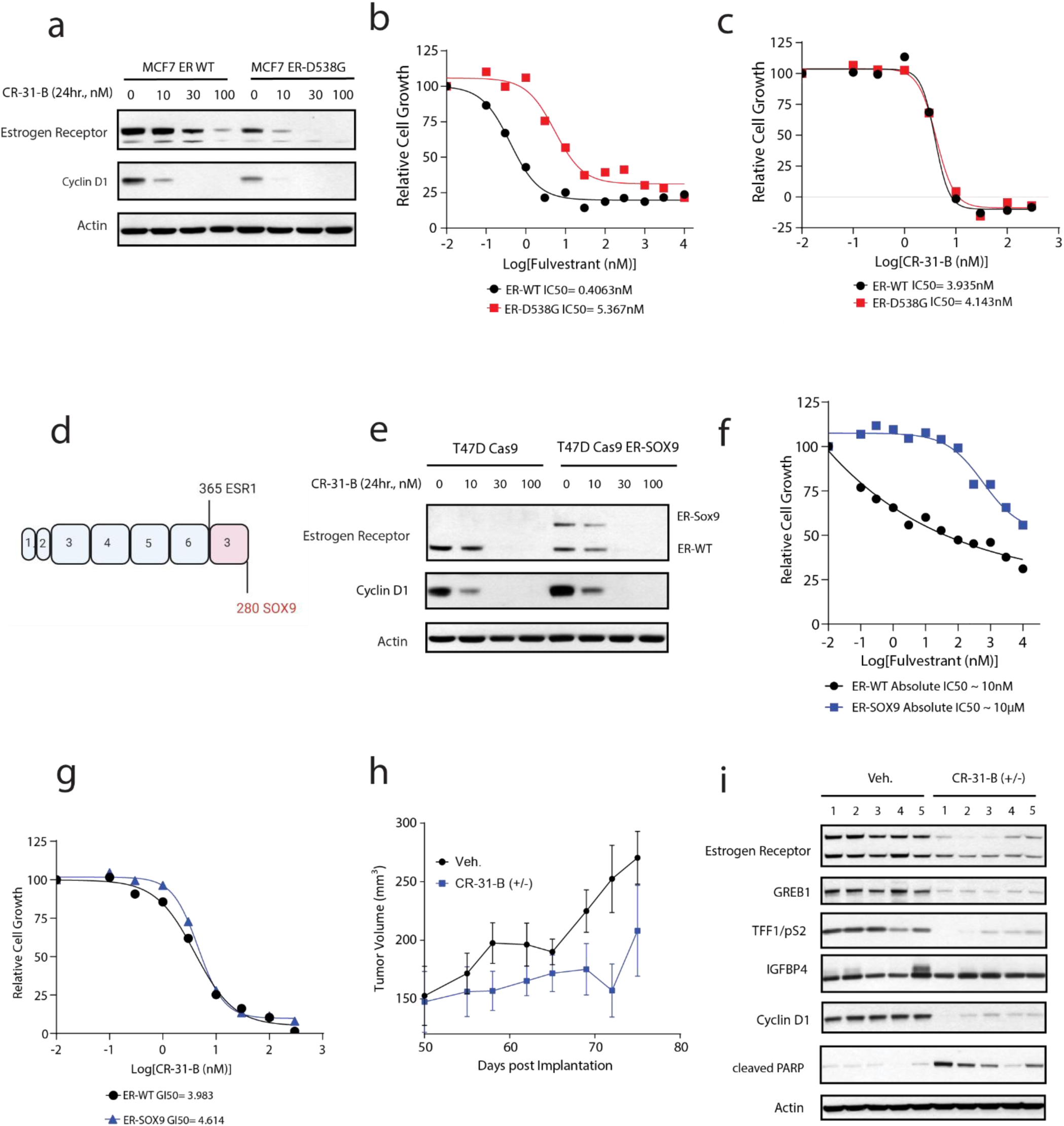
eIF4A inhibition blocks expression of clinically significant ER variants (A) MCF7 expressing either wild type ER or ER-D538G were treated 24hr with increasing doses of CR-31-B (+/-). Data is representative of three independent experiments. (B) MCF7 expressing either wild type ER or ER-D538G were treated 72hr. with increasing doses of fulvestrant. Data is representative of three independent experiments. (C) MCF7 expressing either wild type ER or ER-D538G were treated 72hr. with increasing doses of CR-31-B. Data is representative of three independent experiments. (D) Schematic showing CRISPR Cas9 constructed ESR1-Sox9 fusion. Exons contributed by each protein are indicated. Numbers above and below indicate the residues contributed by each protein (First 365 amino acids of ER and the last 280 of SOX9). (E) T47D Cas9 or T47D Cas9 ESR1-SOX9 were treated for 24Hr with increasing doses of CR-31-B. Data is representative of three independent experiments. (F) T47D Cas9 or T47D Cas9 ESR1-SOX9 were treated for 72hr with increasing doses of Fulvestrant. Data is representative of three independent experiments. (G) T47D Cas9 or T47D Cas9 ESR1-SOX9 were treated for 72hr with increasing doses of CR-31-B. Data is representative of three independent experiments. (H) NSG mice were implanted with estrogen pellets (0.18 mg) for 3d before injection of T47D Cas9 ESR1-Sox9 10million cells/mouse. Once tumors reached 100 mm^3^ mice were treated twice weekly with CR-31-B (+/-) (1mg/kg i.v.). Data is representative of two independent experiments. (I) Immunoblots from xenografts in figure 4G collected 24hr following the final dose of the indicated compounds.

ER fusion proteins have recently been identified in breast cancer, and these have been shown to mediate acquired resistance to hormone receptor antagonists^77,79^. These variants are comprised of the N-terminal portion of wildtype ER fused to a variety of C-terminal partners. The resulting constitutively active fusion protein lacks the hormone binding domain and cannot be inhibited by FDA approved ER inhibitors such as fulvestrant or elacestrant. In order to determine whether the expression of these fusions is sensitive to eIF4A inhibition, we generated a T47D cell line harboring an ESR1-SOX9 fusion previously identified in a human tumor(Figure 4D)^79^. To preserve the untranslated regions of the encoding mRNA, we generated the fusion endogenously using ribonucleofection of Cas9 and guide RNAs against the corresponding introns at the *ESR1* and *Sox9* loci. Confirming estrogen independent growth, cells expressing the ER-SOX9 fusion were enriched after selection in estrogen free media (Figure S4C). Confirming the N-terminal contribution of ER to the fusion, ER could be detected using an N-terminal but not C-terminal targeting antibody against ER (Figure S4C). Treatment with 30nM CR-31-B was able to suppress the ER-SOX9 fusion and cyclin D1 to undetectable levels within 24hr (Figure 4E). The ER-SOX9 fusion was observed to have a similar half-life compared to wildtype ER, with both the ER-SOX9 fusion, as well as wildtype ER being appreciably downregulated at 4hr, and continuing to decrease over the next 20 hours (Figure S4D).

The ER-SOX9 expressing cells were a thousand-fold less sensitive to fulvestrant, with an absolute GI50 of 10µM, vs. 10nM for the wild type ER expressing cells (Figure 4F). ER-SOX9 expressing cells remained sensitive to CR-31-B, exhibiting an GI50 of 4.6nM compared to 3.9nM for ER wildtype cells (Figure 4G). Finally, we tested the effect of eIF4A inhibition on ER-SOX9 expressing xenografts *in vivo*, where Bi weekly doses of 1mg/kg CR-31-B (+/-) produced an inhibition of tumor growth for up to 25 days after treatment (Figure 4H). ER-SOX9 expression and ER targets GREB1, TFF1/pS2, IGFBP4 and cyclin D1 were all suppressed by CR-31-B. Treatment with CR-31-B also induced PARP cleavage, indicating apoptosis initiation in the tumor (Figure 4I). These results show that ER variants with either mutations or deletions in the ligand binding domain that are challenging to treat clinically with anti-estrogen therapy retain their dependence on eIF4A.

### Zotatifin + fulvestrant combination lowers ER expression and Suppresses tumor growth in patients

eIF4A dependent expression of both ER and cell cycle regulators suggests that eIF4A inhibition may be a valid clinical strategy, particularly in combination with standard endocrine therapies like fulvestrant. Based on these results, early phase testing of the eIF4A inhibitor zotatifin (eFT226) includes patients with ER+ metastatic breast cancer (MBC) (NCT04092673)^27^. Similar to other rocaglate eIF4A inhibitors such as silvestrol and CR-31-B, zotatifin binds both eIF4A itself and the 5’UTR of select mRNA^81^ (Figure 5A). Zotatifin is a first in class inhibitor of eIF4A and was designed by enhancing the physicochemical and pharmacokinetic properties of rocaglamide A^81^ (Figure 5A). We observed zotatifin to be qualitatively similar to silvestrol and CR-31-B, with 100nM suppressing both ER and cyclin D1 expression by 24 hours. Zotatifin was ten-fold less potent than silvestrol and CR-31-B at suppressing target expression and cell growth (Figure S5A, S5B).

**Figure 5:**
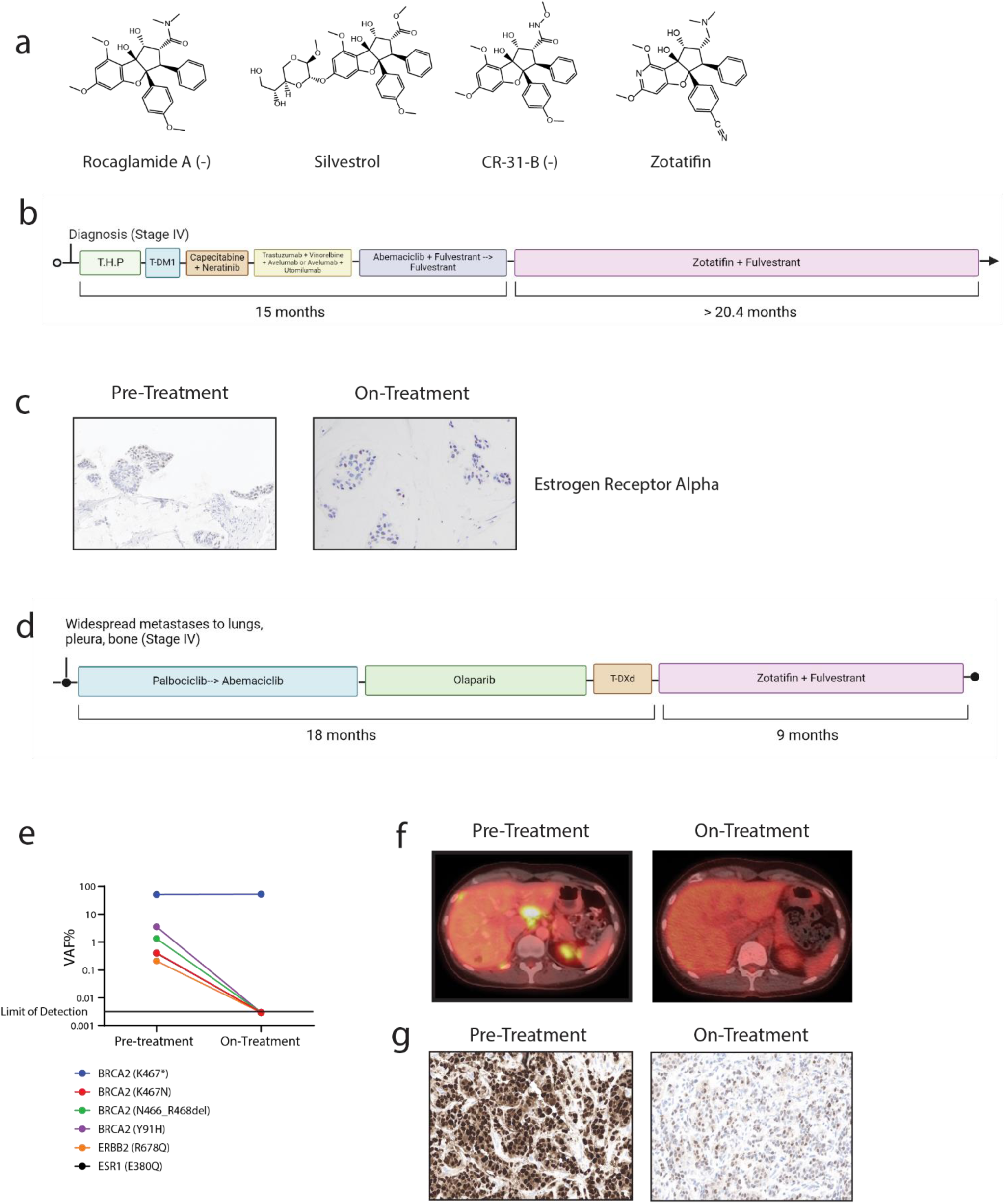
Zotatifin + fulvestrant lowers ER expression and suppresses tumor growth in patients (A) Chemical Structures of Rocaglate family inhibitors of EIF4A including clinical compound Zotatifin. (B) Treatment timeline for patient 1, depicting relative times on various treatments including zotatifin + fulvestrant. T.H.P= docetaxel, trastuzumab, and pertuzumab, TDM1= Kadcyla (ado-trastuzumab emtansine) (C) IHC staining for Estrogen Receptor alpha pre and post treatment with Zotatifin + Fulvestrant. (D) Treatment timeline for patient 2, depicting relative times on various treatments including zotatifin + fulvestrant. TDXD= Trastuzumab Deruxtecan (E) Somatic Cell free DNA pre and post-treatment with zotatifin + fulvestrant (F) PET CT showing pre and post-treatment with zotatifin + fulvestrant (G) IHC staining for Estrogen Receptor alpha pre and post treatment with Zotatifin + Fulvestrant.

The ER+ metastatic breast cancer dose expansion cohort currently features two treatment arms: zotatifin + fulvestrant (ZF) and zotatifin + fulvestrant + abemaciclib (ZFA). Clinical responses have been seen in heavily pretreated MBC patients in both the ZF and ZFA cohorts^27^. Notably all patients on trial have had previous exposure to and developed resistance to fulvestrant and/or CDK4/6 inhibitors, suggesting that zotatifin both adds to combined endocrine + CDK4/6 inhibitor therapy and may potentially re-sensitize patients in the case of acquired resistance. Both ZF and ZFA regimens were well tolerated, with no dose limiting toxicities or grade 5 adverse events^27^.

Results thus far corroborate the mechanistic findings detailing the interaction between eIF4A inhibition and ER downregulation as a potential driver of clinical response. One patient treated on the ZF doublet arm (initially with ER+PR+HER2+ disease, subsequently with loss of HER2 positivity, likely with some degree of intratumoral HER2 heterogeneity) enrolled on the trial as 6th line therapy for progressive metastatic disease and experienced disease stability for over 19 months and continues on treatment (Figure 5B). Analysis of pre-treatment biopsies demonstrated ER expression by immunohistochemistry of 40%, while on-treatment biopsies showed near complete suppression of ER, with expression decreased to <10%. (Figure 5C).

A second patient in the ZF doublet arm enrolled on trial as 4^th^ line therapy for progressive metastatic disease after receiving endocrine therapy + CDK4/6 inhibition, a PARP inhibitor, and the ADC trastuzumab deruxtecan in the metastatic setting(Figure 5D). This patient showed a brisk RECIST-confirmed partial response (-55%) at 8 weeks, with regression of chest wall and liver lesions, as well as a 100% decline in somatic ctDNA 6 weeks into treatment (Figure 5E, 5F). CtDNA analysis revealed a ESR1 E380Q^82^ mutation and a somatic activating ERBB2 R678Q mutation, thereby providing proof of principle that these endocrine therapy-resistant subclones were sensitive to combination zotatifin therapy (Figure 5E). While ER suppression at the protein level was not as complete as that seen in the first patient, pre-treatment and on-treatment biopsies showed a significant change from >99% ER expression down to 60% (Figure 5G). Overall, these data showing clinical responses coupled with this drug’s well-tolerated^27^ side effect profile demonstrates that combination zotatifin therapy may represent a clinically viable strategy for treating metastatic ER+ breast cancer.

## Discussion

In this work, we demonstrate that estrogen receptor alpha (ER) translation is dependent on eIF4A and that targeting eIF4A can potently inhibit ER expression and promote antitumor effects in ER+ breast cancers. These findings open up a novel therapeutic opportunity for selectively targeting the translation of a few key modulators of tumor growth in the context of estrogen receptor driven breast cancer.

Canonical eukaryotic translation is controlled by the m7G cap binding protein, eIF4E. By virtue of mTORC1 inactivation, eIF4E availability and hence translation is suppressed when nutrients (glucose, amino acids etc.) are limited. However, evolution has selected for a subset of mRNAs whose translation can proceed non-canonically (cap-independently) and are therefore not tied to nutrient status. The logic being that developmentally essential or stress response genes can be translated during fasting or starvation. Non-canonical translation is an area of active investigation with many eIF4E independent mechanisms described^18,20–23,47,48,83–95^. Among the transcripts capable of eIF4E independent translation are many oncogenes including Myc^60,96–99^, Bcl-2^100,101^, c-jun^20,102^, and developmentally essential genes among the Hox clusters^103,104^. Here we demonstrate estrogen receptor alpha (*ESR1*) is another such gene. Frequently, the elements mediating cap-independent translation are present in the transcript 5’ untranslated region (5’ UTR). Such elements are broadly termed, IRESes (Internal ribosome entry sites) and often serve as binding sites for selectively employed initiation factors or to allow the ribosome to link directly with the mRNA transcript. IRESes often take on complex 3D structures which require unwinding and remodeling during initiation, and eIF4A is one such protein employed for this purpose.

G-quadraplex elements are one 3D structure that have been implicated in both cap-independent translation and in conferring eIF4A dependence to select mRNAs^59,87^. These elements are thermodynamically stable 3D arrangements of guanosines conjugating a monovalent cation^105^. Indeed, the 5’ UTR of ESR1 transcript variant 1 (NM_000125.4) is predicted to contain an abundance of overlapping and non-overlapping G-quadraplex elements^106^. Silvestrol, CR-31-B and zotatifin have all been shown to block eIF4A dependent unwinding of these structures^24,96^ Here we demonstrate numerous potential clinical applications of eIF4A dependent ER regulation. We show that suppressing ER expression in this manner reduces ER target gene expression and is synergistic with fulvestrant in blocking ER functionality and in blocking cell growth. eIF4A inhibitors can also suppress expression of clinically significant ER mutants and difficult to treat ER fusions. Analogously, we have also observed that eIF4A inhibition can suppress androgen receptor translation (AR) as well as the AR variant 7 (AR-V7) which lacks the ligand binding domain and signals in an androgen independent manner (data not shown).

Supported by our mechanistic findings, we show clinical data from two patients where combination zotatifin and endocrine therapy was clinically active. These patients both show clear reduction in ER at the protein level, and suppressed ER expression may be a useful marker of zotatifin efficacy in patients going forward. This drug’s activity in these heavily pretreated patients may also stem from the fact that it targets the G1/S checkpoint through additional key nodes such as cyclin D and CDK4. Because eIF4A inhibitors lower both ER and cyclin D/CDK4 levels, we reason that eIF4A inhibitors can concurrently enhance the activity of both endocrine therapy and cell cycle targeting agents.

This clinical trial is ongoing and continues to accrue, and embedded within this trial is a significant translational component where the above questions continue to be investigated, as biomarkers of sensitivity to zotatifin will be key to the ultimate success of this clinical program. In general eIF4A inhibition represents a novel strategy by which ER+ metastatic breast cancers can now be targeted in the clinic.

## Supporting information

eIF4A controls translation of estrogen receptor alpha and is a therapeutic target in advanced breast cancer supplement

Table S1 Proteomic changes in MCF7 24hr following silvestrol treatment

## Acknowledgements

We are grateful to all members of the Rosen Lab past and present for helpful discussions and advice. Thanks to Ventura lab for help with CRISPR experiments. NCI Core Grant P30 CA008748 is gratefully acknowledged for partial funding of both the Organic Synthesis and Mouse Pharmacology Core Facilities at MSKCC. The Organic Synthesis Core is also partially funded through NCI grant R50 CA243895. F. Pareja is funded in part by an NIH/NCI P50 CA24779 01 grant and by a Starr Cancer Consortium Grant. The Wendel lab gratefully acknowledges funding from R35 CA252982, Harrington Discovery Institute’s 2022 Scholar Innovator Award, Starr Technology Commercialization Fund (Starr TCF) and P30 CA008748. S. Chandarlapaty is supported by NIH Cancer Center Support Grant P30-CA008748 and NIH R01CA245069 and the BCRF. This research was supported by grants (to N.R.)(and long term support from the BCRF) from the National Institutes of Health (NIH) P01-CA129243; R35 CA210085; the Geoffrey Beene Cancer Research Center; the Emerson Collective Research Grant, The NIH MSKCC Cancer Center Core Grant P30 CA008748 and Experimental Therapeutics Center. All illustrations were designed with BioRender and all chemical structures were designed with ChemDraw.

## Author Contributions

J.A.B., N.R., and S.C. conceived the hypotheses. J.A.B., S.C. and N.R. wrote the manuscript. J.A.B, M.S., M.D., E.Y.R., N.M., C.A., S.G., R.K., J.X., M.M, Z.L. E.S., S.W., D.W., S.S., G.G.C., P.A.G., G.Y., O.O., H.-G.W, S.C. and N.R. contributed to experimental design and analysis. J.A.B., M.S., M.D., C.A., S.G., R.K., J.M.R., X.C., M.M., Q.C. and F.P performed experiments, and E.Y.R., N.M., and F.P. performed the clinical research.

## METHODS

### Cell Lines

All cell lines were purchased from American Type Culture Collection (ATCC). MCF7 (HTB-22), T47D (HTB-133), ZR-75-1 (CRL-1500), BT474 (HTB-20), SKOV3 (HTB-77), T47D-Kbluc (CRL-2865). All Cell lines were maintained in DMEM/F12 supplemented with 10% Fetal Bovine Serum (FBS) and 1% penicillin and streptomycin. Stably generated cell lines including those expressing rtTA3 and tetracycline inducible constructs were maintained in DMEM/F12, 10% Tetracycline free Fetal Bovine Serum (Tet-free FBS), and 1% penicillin and streptomycin. All Cells were maintained in a humidified incubator with 5% CO2 at 37 °C.

### Immunoblotting

Cells were collected in ice cold PBS and lysed with RIPA lysis buffer (Pierce #89901) supplemented with Halt protease and phosphatase inhibitors (Pierce Chemical). Lysates were briefly sonicated before centrifugation at 20,000 × g for 5 minutes at 4°C. The supernatant was collected, and protein concentration was determined using the BCA kit (Pierce) per manufacturer’s instructions. Equal amounts of protein (20μg) in cell lysates were separated by SDS–PAGE, transferred to nitrocellulose membranes (GE healthcare), immunoblotted with specific primary and secondary antibodies and detected by chemiluminescence with the ECL detection reagents from Thermo Fisher or Millipore. Protein quantification when applicable was performed using Fiji.

### Antibodies

Anti-puromycin (Kerafast EQ0001), pAKT S473 (Cell Signaling Technology (CST) 4060), p4EBP1 T37/46 (CST 2855), Cyclin D1 (CST 55506), Estrogen Receptor Alpha N-terminus (CST 13258), Estrogen Receptor Alpha C-terminus (CST 8644), Progesterone Receptor (CST 8757), EIF4G (CST 2498), EIF4E (CST 9742), 4EBP1 (CST 9644), Beta Actin (CST 4967), Myc (CST 18583), GATA3 (CST 5852), Cyclin D3 (CST 2936), CDK4 (CST 12790), pRb S780 (CST 9307), E2F1 (CST 3742), GREB1 (CST 65171), TFF1/pS2 (CST 15571), IGFBP4 (CST 31025), cleaved PARP (CST 5625). Secondary Goat anti-Rabbit IgG (H+L) Secondary Antibody HRP (Thermo Fisher 65-6120).

### Plasmids

All plasmids for the Dual Luciferase Reporter were generated by subcloning gene body (GAPDH) or 5’UTR elements (ER, Myc) into Addgene 45642, pLenti CMV rtTA3 Hygro (w785-1) (Addgene: 26730), pCW57.1-4EBP1_4xAla (Addgene: 38240).

### Generation of tetracycline inducible cell lines

293GP cells were plated in 10cm plates and transfected using lipofectamine 2000 with 6μg retroviral and 1.5μg pMD2G plasmids. Supernatent was harvested from packaging cells on two consecutive days beginning 24hr following transfection. 8μg/ml polybrene was added to virus containing supernatant before adding to target cells. MCF7 parental cells were first infected with a CMV driven expression construct encoding rtTA3 (addgene:26730). Cells were selected in 250μg/ml hygromycin for 7 days. 293T cells were plated in 10cm plates and transfected using lipofectamine 2000 with 6μg lentiviral pCW57.1-4EBP1_4xAla (Addgene: 38240), 1.5μg pMD2G and 4.5μg psPax2. Virus containing supernatant with 8μg/ml polybrene was collected on two consecutive days and added to MCF7-rtTA3. Cells were selected for 3 days in 2μg/ml puromycin. Cells were thereafter maintained in DMEM F12 containing tetracycline-free FBS and 1% Penicillin/Streptomycin supplemented with 0.5ug/ml puromycin and 100ug/ml hygromycin.

### In vitro cap-binding affinity assay

Experiments were conducted using the protocol from^11^. 200μg of lysate was incubated with m7G conjugated agarose beads (Jena Biosciences) for 2hr at 4 degrees with rotation. Beads were washed 3 times with ice cold lysis buffer. Bound proteins were eluted with 1x loading buffer with heating at 95 degrees for 5 min. EIF4F complex composition was analyzed by immunoblotting using indicated antibodies.

### AHA labeling and click chemistry

MCF7 were starved of methionine for 30min with simultaneous administration of indicated treatments, then pulsed with 100μM AHA (Thermo Fisher: C10102) for 2hr. 200μg of protein was used for click chemistry and was performed using biotin-alkyne and protein reaction buffer kits (Thermo Fisher: C10276). AHA-biotin-alkyne labeled proteins were pulled down with streptavidin beads and the indicated proteins were analyzed by immunoblotting.

### Dual Luciferase Reporter

5’ UTR elements or part of the GAPDH gene body^102^ were cloned into the dual luciferase assay construct (Addgene:45642). 1.5 million cells/6cm plate were transfected with 2μg of the construct using lipofectamine 2000 at a ratio of 3:1 lipofectamine to μg DNA. Firefly and Renilla luciferase activity was measured 24hr post transfection via dual luciferase assay reporter system (Promega) according to the manufacturer’s instructions.

### mRNA extraction and RT-qPCR

mRNA was isolated using Trizol based phenol chloroform extraction. cDNA was synthesized using Quantitect Reverse Transcription kit (Qiagen). Transcript quantification was done using Applied Biosystems Taqman probes and ABI 7500 real-time quantitative PCR system. For data analysis, cycle numbers were normalized to housekeeping gene, Rplp0, and then to untreated control (2^−ΔΔ*C*t^).

### Chromatin Immunoprecipitation Assay (ChIP)

MCF7 were plated in normal DMEM F12. Cells were then washed twice and media was changed to DMEM F12 lacking phenol red and containing charcoal stripped FBS (-E2), with or without indicated drug (s) for an additional 24hr. Cells were then stimulated with 10nM Estradiol for 1hr. ChIP was performed using the Simple ChIP Enzymatic Chromatin IP kit (Agarose Beads) from Cell signaling (#9002) according to the manufacurer’s instructions. ER enhancer binding was measured via PCR amplification of the ER enhancer upstream of TFF1/pS2. Primers were from Cell Signaling (#9702)

### Quantification of Cell Growth and Viability

Cells were seeded into 96-well plates at between 2000-5000 cells per well. Cell growth was quantified using the ATP-Glo assay (Promega, G7572). For each condition at least 3 replicates were measured. For GI50 curves, cells were treated for 72hr and day 0 values were subtracted from each group. Sigmoidal growth inhibition curves were calculated using a four-parameter model in Graph Pad Prism 8.

### LC–MS proteomic analysis

Cells were lysed in 8 M Urea, 200 mM EPPS (4-(2-Hydroxyethyl)-1-piperazinepropanesulfonic acid), pH 8.5 with protease (complete mini EDTA-free, Roche) and phosphatase inhibitors (cocktail 2 and 3, Sigma). Samples were then sonicate for 1 minute and a BCA assay was used to determine the protein concentrations. Aliquots of 100 µg were taken for each sample (based on BCA assay) and reduced with 5 mM TCEP (tris(2-carboxyethyl) phosphine hydrochloride), alkylated with 10 mM IAA (iodoacetamide), and quenched with 10 mM DTT (dithiothreitol). Samples were diluted to 100 µL with lysis buffer and precipitated by chloroform–methanol. Pellets were resuspended in 50 µL 200 mM EPPS buffer, digested with Lys-C protease at a 1:50 protease-to-protein ratio for 4 hrs at 37 °C, then overnight with trypsin (1:50) at 37 °C. Anhydrous acetonitrile was added at a final volume of 30%. TMT (11-plex) reagents were added to peptides at a 2:1 (TMT reagent-to-peptide ratio) and incubated for 1 h at room temperature. A label check was performed to determine mixing ratios, labelling efficiency, and number of missed cleavages by pooling 1 µL from each sample, desalting, then analyzing by mass spectrometry. Samples were mixed 1:1 across all channels, dried to remove acetonitrile, then desalted using C18 solid-phase extraction (SPE) Sep-Pak (Waters), and vacuum centrifuged to dryness. Dried samples were reconstituted in 1 mL of 2% ACN/25 mM ABC. Peptides were fractionated into 48 fractions. An Ultimate 3000 HPLC (Dionex) coupled to an Ultimate 3000 Fraction Collector using a Waters XBridge BEH130 C18 column (3.5 um 4.6 × 250 mm) was operated at 1 mL/min. Buffer A consisted of 100% water, buffer B consisted of 100% acetonitrile, and buffer C consisted of 25 mM ABC. The fractionation gradient operated as follows: 1% B to 5% B in 1 min, 5% B to 35% B in 61 min, 35% B to 60% B in 5 min, 60% B to 70% B in 3 min, 70% B to 1% B in 10 min, with 10% C the entire gradient to maintain pH. The 48 fractions were then concatenated to 12 fractions (i.e., fractions 1, 13, 25, 37 were pooled, followed by fractions 2, 14, 26, 38, etc.) so that every 12th fraction was used to pool. Pooled fractions were vacuum-centrifuged then reconstituted in 1% ACN/0.1% FA for LC–MS/MS. Fractions were analyzed by LC-MS/MS using a Thermo Easy-nLC 1200 (Thermo Fisher Scientific) with a 50 cm (inner diameter 75µm) EASY-Spray Column (PepMap RSLC, C18, 2µm, 100Å) heated to 60°C coupled to a Orbitrap Fusion Lumos Tribrid Mass Spectrometer (Thermo Fisher Scientific). Peptides were separated at a flow rate of 300nL/min using a linear gradient of 1 to 30% acetonitrile (0.1% FA) in water (0.1% FA) over 4 hrs and analyzed by SPS-MS3. MS1 scans were acquired over a range of m/z 375–1500, 120K resolution, AGC target of 4 × 105, and maximum IT of 50 ms. MS2 scans were acquired on MS1 scans of charge 2–7 using isolation of 0.7m/z, collision-induced dissociation with activation of 35%, turbo scan and max IT of 50 ms. MS3 scans were acquired using specific precursor selection (SPS) of 10 isolation notches, m/z range 100–1000, 50K resolution AGC target of 1e5.

### TMT data analysis

Raw data files were processed using Proteome Discoverer (PD) version 2.4.1.15 (Thermo Scientific). For each of the TMT experiments, raw files from all fractions were merged and searched with the SEQUEST HT search engine with a Homo sapiens UniProt protein database downloaded on 2019/01/09 (176,945 entries). Methionine oxidation was set as variable modification, while cysteine carbamidomethylation, TMT6plex (K), and TMT6plex (N-term) were specified as fixed modifications. The precursor and fragment mass tolerances were 10 ppm and 0.6 Da, respectively. A maximum of two trypsin missed cleavages were permitted. Searches used a reversed sequence decoy strategy to control peptide false discovery rate (FDR) and 1% FDR was set as the threshold for identification.

### ER Reporter Assay

T47D KBluc were plated in normal DMEM F12. Cells were then washed twice and media was changed to DMEM F12 lacking phenol red and containing charcoal stripped FBS (-E2), with or without indicated drug (s) for an additional 24hr. Cells were then stimulated with estradiol for a final 24hr. Firefly luciferase activity was measured via dual luciferase assay reporter system (Promega) according to the manufacturer’s instructions.

### In Vivo Tumor Models

Eight-week-old athymic nu/nu female mice (MCF7) (Harlan Laboratories), or NOD scid gamma mice (T47D) were injected subcutaneously with 10 million cells together with matrigel (BD Biosciences). 17β-Estradiol pellets (0.18 mg or 0.72mg/90 days release) (Innovative Research of America) were implanted subcutaneously 3 days before tumor cell inoculation. Once tumors reached an average volume of 100 mm^3^, mice were randomized (*n* = 3-5 mice per group) to receive CR-31-B(+/-) in 10% Captisol 1mg/kg i.v. twice/week, Fulvestrant in 5% EtOH and 95% castor oil twice/week. Tumors were measured twice weekly using calipers, and tumor volume was calculated using the formula: length × width^2^ × 0.52. All tumors were collected 24hr following the final dose. Samples were lysed and processed as previously described^107^.

### EdU Labeling and Cell Cycle Analysis

MCF-7 that had been treated for 48 with fulvestrant, silvestrol or fulvestrant and silvestrol were incubated with EdU (10μM) for 1.5h at 37C. Cells were then processed with a Click-iT Plus EdU Alexa Fluor 594 flow cytometry kit (Thermo Fisher, C10646) following the manufacturer recommendation. Cells were analyzed by flow cytometry on a LSRFortessa instrument (BD Biosciences) and data were analyzed using FlowJo (TreeStar).

### Generation of ER-SOX9 Fusion

Cas9 guide RNA ribonucleoprotein complex was used to generate ESR1-SOX9 fusion. Human guide RNAs targeting human ESR1 intron 6 (GCTCCTGAACGAATACACTG) or human SOX9 intron2 (CGGGACGGAGATAGCTTGTC) were designed as crRNAs (Alt-R CRISPR-Cas9 crRNA) and synthesized by Integrated DNA Technologies. The RNP complex was assembled according to manufacturer’s instructions. Briefly, guide RNA was assembled by mixing equimolar amounts of crRNA and tracrRNA (Alt-R CRISPR-Cas9 tracrRNA-ATTO-550) and heating to 95°C for 5 min. The crRNA-tracrRNA mixture was annealed by slow cooling to room temperature. Cas9 (Alt-R S.p. Cas9 Nuclease V3) and gRNA were incubated for 10 min at room temperature to allow RNP formation. RNP complex was delivered into 1x10^5 T47D cells by 4D-Amaxa Nucleofector System (Lonza) using the SE Cell Line 4D Nucleofector X Kit (Lonza) and program EN-104 was used for nucleofection. 48 hours post nucleofection, genomic DNA was isolated using phenol:chloroform:isoamyl alcohol (Invitrogen). To amplify ESR1-SOX9 genomic fusion, forward ESR1 primer (GAGGTGAGAGGATGGCTTGA) and reverse SOX9 primer (TAGCTGCCCGTGTAGGTGAC) were employed to uniquely amplify the ESR1-SOX9 fusion. For reverse transcription-PCR, total RNA was isolated from cells using Trizol reagent (Invitrogen) and cDNA was generated using SuperScript III First-Strand Synthesis System (Invitrogen). To amplify the chimeric RNA transcript, forward primer against ESR1 (GACAGGGAGCTGGTTCACAT) part of the chimeric transcript and reverse primer (ATCGAAGGTCTCGATGTTGG) against SOX9 was used. All PCR products were confirmed by sanger sequencing. All crRNAs and primer sequences above are listed in 5’-3’.

Study of eFT226 in Subjects With Selected Advanced Solid Tumor Malignancies (Zotatifin) This is a US, open-label, phase 1-2 Dose-Escalation and Cohort-Expansion Study of Intravenous Zotatifin (eFT226) in Subjects With Selected Advanced Solid Tumor Malignancies (NCT04092673). The study was conducted in accordance with the Declaration of Helsinki and was reviewed and approved by the institutional review board. Written informed consent was obtained from all patients before study entry. Patients eligible for study participation were ≥18 years old, had histological or cytological confirmation of breast cancer, metastatic disease or locoregionally recurrent disease which is refractory or intolerant to existing therapy(ies) known to provide clinical benefit, and prior treatment had included a CDK4/6 inhibitor. Tumor is ER+ (defined as ER IHC staining > 0%). Adverse events were graded per Common Terminology Criteria for Adverse Events (CTCAE) 5.0, and responses were evaluated per RECIST version 1.1. Levels of ctDNA in plasma were assessed by next-generation sequencing, using 74-gene Guardant360. Immunohistochemical (IHC) staining for Estrogen receptor (ER) was conducted using the Leica Bond-3 automated stainer platform (Leica, Buffalo Grove, IL). Following antigen retrieval using a citrate-based pH 6 epitope retrieval solution, four-micron formalin fixed paraffin embedded (FFPE) tissue sections were incubated with the primary anti-ER 6F11 antibody (Leica) for 20 minutes. A polymeric kit (Refine, Leica) was used as secondary reagent for detection. ER status was evaluated by IHC by a board-certified pathologist (FP) following the American Society of Clinical Oncology/College of American Pathologists (ASCO/CAP) guidelines (PMID: 20404251), considering 1% of positive tumor nuclei as threshold for ER positivity. Data cutoff for the presented results is January, 2024.

### Statistical Analysis

The details of statistical analysis of experiments can be found in the figure legends. All data were plotted as mean +/- standard deviation with the exception of Figures: 3D, 3F and 4H which were plotted as mean +/- SEM. Statistical analysis of differences between two groups was performed using two-tailed Student’s t-test, and p < 0.05 was defined as significant. One-way ANOVA analysis was performed to compare the means of more than two groups. All analysis was conducted using GraphPad Prism 8.0. Independent experiments were conducted with a minimum of two biological replicates per condition to allow for statistical comparison.

