## Supplementary material for "eIF4A controls translation of estrogen receptor alpha and is a therapeutic target in advanced breast cancer": eIF4A controls translation of estrogen receptor alpha and is a therapeutic target in advanced breast cancer supplement

### Supplemental Figure 1

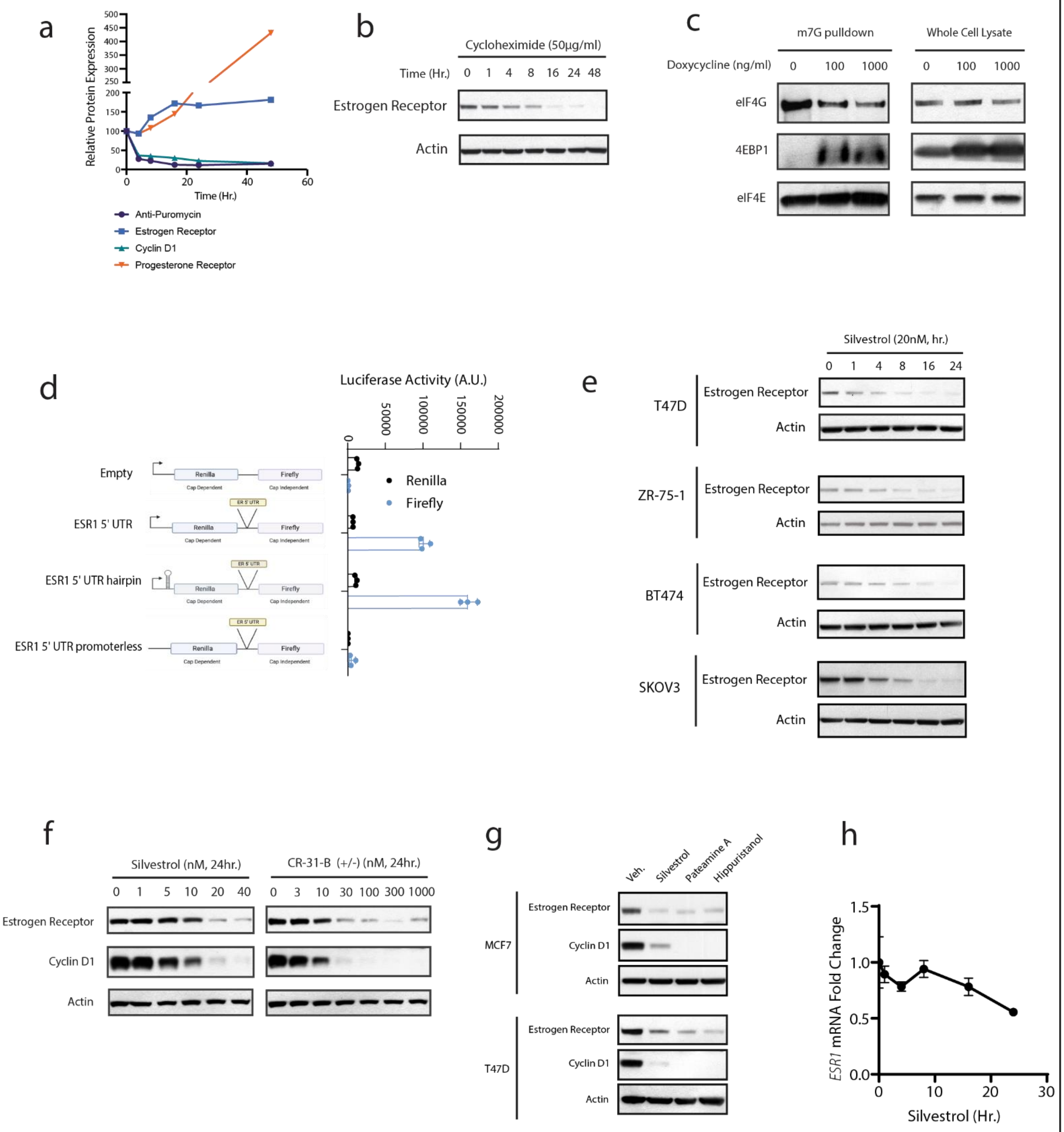

Supplemental Figure 1

(A) Protein expression and puromycin incorporation from Fig 1B quantified using densitometry.

(B) MCF7 cells were treated for time t with cycloheximide (50μg/ml). Data is representative of three independent experiments.

(C) MCF7 were treated 24hr with increasing doses of doxycycline to induce 4EBP1 (T37A/T46A/S65A/T70A) "4EBP1 4A" expression. Cell lysates were then used for immunoblotting (Whole Cell Lysate) or m7G cap-pulldowns to assess EIF4G and 4EBP1 binding to EIF4E. Data is representative of three independent experiments.

(D) MCF7 were transfected with dual luciferase constructs, containing an empty cassette or the ESR1 variant1 5'UTR. The 5' UTR of ESR1 was subcloned into vectors containing a hairpin upstream of the renilla cassette, or lacking any upstream promoter. n=3 samples per group. Data is representative of three independent experiments.

(E) Indicated ER+ cell lines were treated for time t with 20nM silvestrol. Immunoblots were performed to assess ER expression as a function of time. Data is representative of three independent experiments.

(F) MCF7 cells were treated for 24hr with inc. doses of eIF4A inhibitors, silvestrol or CR-31-B (+/-). Data is representative of three independent experiments.

(G) MCF7 or T47D were treated 24hr with EIF4A inhibitors, Silvestrol (20nM), Pateamine A (1µM) or Hippuristanol (1µM). Data is representative of three independent experiments.

(H) MCF7 were treated for indicated times with 20nM Silvestrol and ESR1 mRNA was measured via qRT-PCR. Data is representative of three independent experiments.

Supplemental Figure 2

a

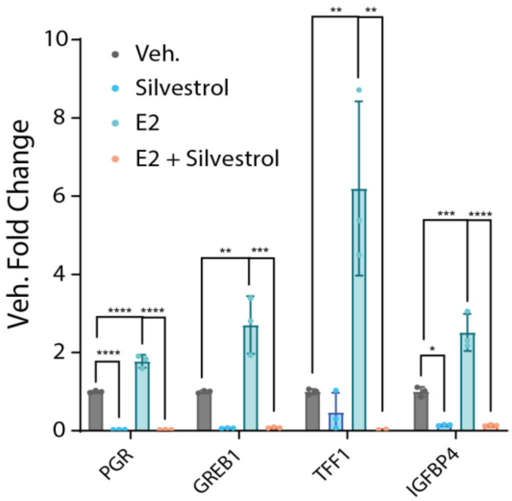

Supplemental Figure 2

(A) T47D were plated in DMEM F12 containing charcoal stripped FBS and lacking phenol red, followed by treatment with silvestrol (20nM) for 24hr. Cells were then stimulated with 10nM estradiol for an additional 24hr. ER target gene expression was analyzed via RT-qPCR. P-values were determined by one-way ANOVA for each gene. P<.05 depicted as *, P<.01 depicted as **, P< .001 depicted as *** and p< .0001 as ****. Data is representative of three independent experiments.

#### Supplemental Figure 3

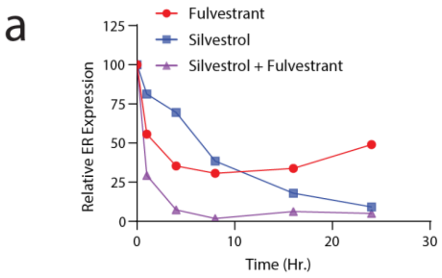


| | Fulvestrant | Silvestrol | Silvestrol + Fulvestrant |
|---|---|---|---|
| AUC | 935.7 | 867.2 | 215.2 |

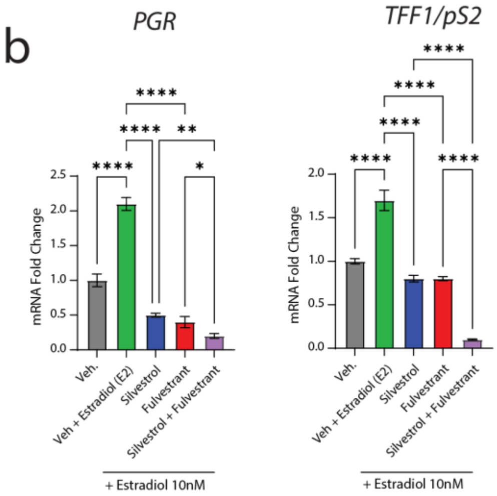


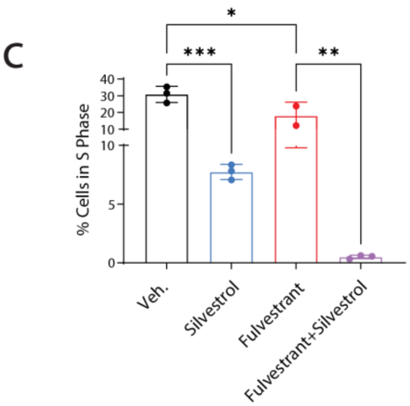


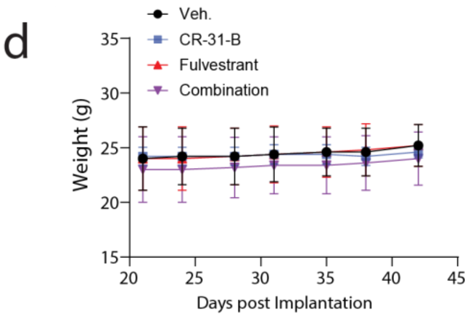

Supplemental Figure 3

(A) Quantification of protein expression from Figure 3A. Densitometry was performed using Fiji (Image J).

(B) MCF7 were plated in phenol red free charcoal stripped FBS DMEM F12. Cells were then treated with Veh. or Silvestrol (5nM), Fulvestrant (3nM) or the combo for 24hr. Cells were finally stimulated with 10nM estradiol for an additional one day before isolation of mRNA for RT-qPCR. P-values were determined by one way ANOVA for each gene. P<.05 depicted as *, P<.01 depicted as **, P< .001 depicted as *** and p< .0001 as ****. Data is representative of three independent experiments.

(C) Percentage of cells in S phase as measured by 1.5h EdU pulse incorporation of MCF-7 treated for 48h with fulvestrant (3nM), silvestrol (5nM) or the combination. Mean of three biological replicates is plotted. P-values were determined by one way ANOVA. P<.05 depicted as *, P<.01 depicted as **, and P< .001 depicted as ***. Data is representative of two independent experiments.

(D) Kinetic measurement of mouse weights over the course of the "high estrogen experiment". Data is representative of three independent experiments. Data is representative of two independent experiments.

Supplemental Figure 4

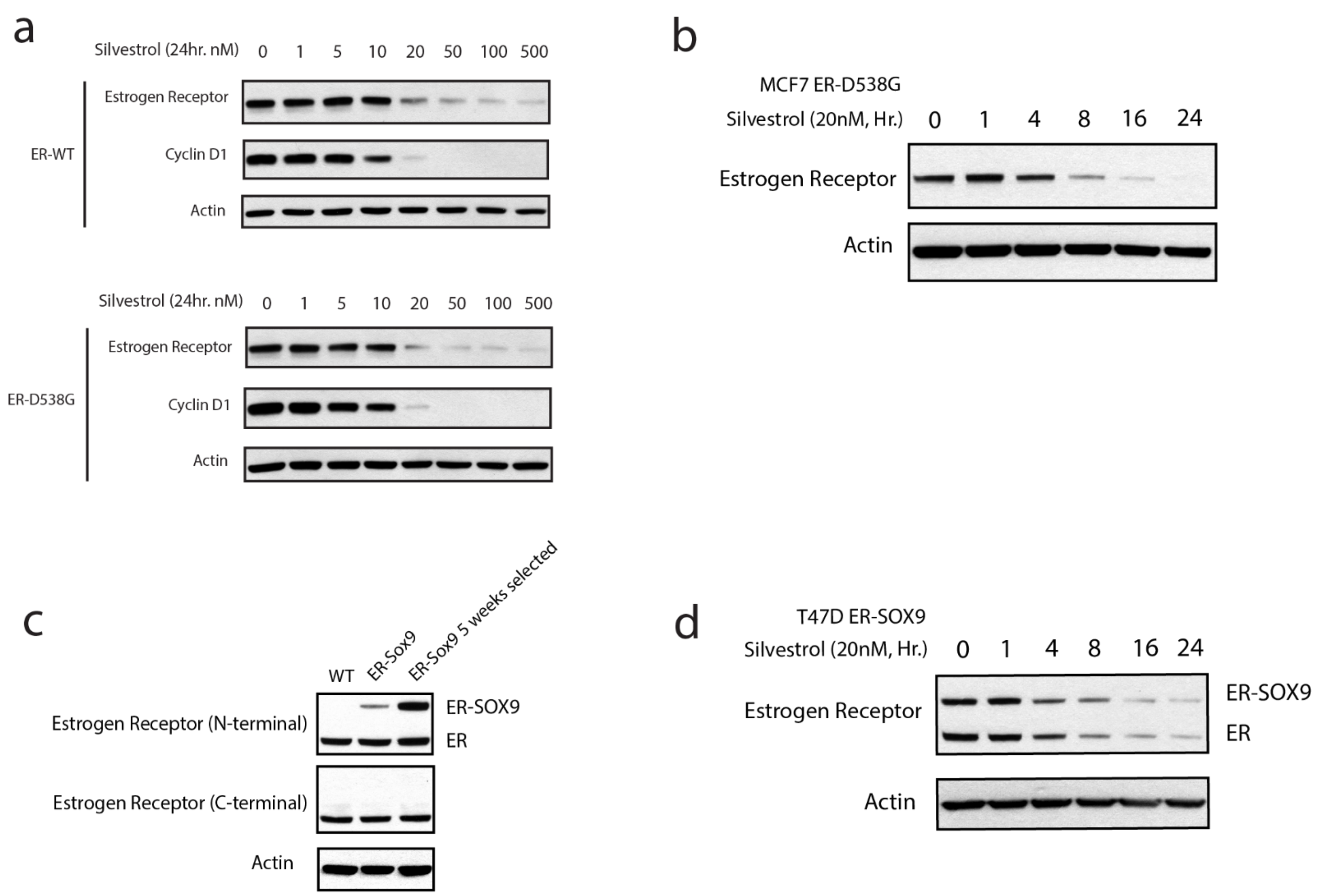

Supplemental Figure 4

(A) MCF7 expressing either wild type ER or ER-D538G were treated 24hr with increasing doses of silvestrol. Data is representative of three independent experiments.

(B) MCF7 ER-D538G expressing cells were treated with 20nM silvestrol for the indicated times. Data is representative of three independent experiments.

(C) Immunoblot showing expression of ER-SOX9 fusion protein in cells selected in estrogen free conditions for 5 weeks.

(D) T47D Cas9 ER-SOX9 treated for the indicated times with 20nM Silvestrol. Data is representative of three independent experiments.

Supplemental Figure 5

a

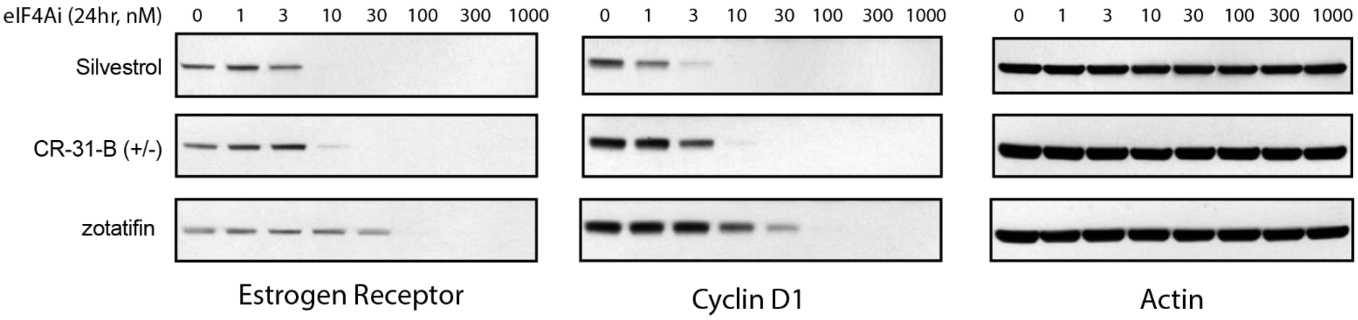


b

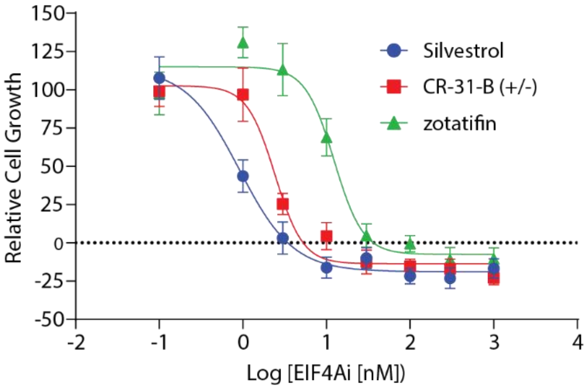

Supplemental Figure 5

(A) T47D cells were treated for 24 hours with increasing doses of the indicated eIF4A inhibitor. Data is representative of three independent experiments.

(B) T47D cells were treated for 72 hours with increasing doses of the indicated eIF4A inhibitor. Data is representative of three independent experiments. Relative cell growth was quantified using ATP-glo.
